# Liquid-Liquid Phase Separation of Patchy Particles Illuminates Diverse Effects of Regulatory Components on Protein Droplet Formation

**DOI:** 10.1101/294058

**Authors:** Valery Nguemaha, Huan-Xiang Zhou

**Affiliations:** Department of Physics and Institute of Molecular Biophysics, Florida State University, Tallahassee, FL 32306, United State; Department of Chemistry and Department of Physics, University of Illinois at Chicago, Chicago, IL 60607, United States

## Abstract

Recently many cellular functions have been associated with membraneless organelles, or protein droplets, formed by liquid-liquid phase separation (LLPS). Proteins in these droplets often contain RNA-binding domains, but the effects of RNA on LLPS have been controversial. To gain better understanding on the roles of RNA, here we used Gibbs-ensemble simulations to determine phase diagrams of two-component patchy particles, as models for mixtures of proteins with RNA or other regulatory components. Protein-like particles have four patches, with attraction strength *ε*_PP_; regulatory particles experience mutual steric repulsion but have two attractive patches toward proteins, with the strength *ε*_PR_ tunable. At low *ε*_PR_, the regulator, due to steric repulsion, preferentially partitions in the dispersed phase, thereby displacing the protein into the droplet phase and promoting LLPS. At moderate *ε*_PR_, the regulator starts to partition and displace the protein in the droplet phase, but only to weaken bonding networks and thereby suppress LLPS. At *ε*_PR_ > *ε*_PP_, the enhanced bonding ability of the regulator initially promotes LLPS, but at higher amounts, the resulting displacement of the protein suppresses LLPS. These results illustrate how RNA can have disparate effects on LLPS, thus able to perform diverse functions in different organelles.

## Introduction

There is intense current interest in membraneless organelles, thought to be formed by liquid-liquid phase separation (LLPS) of protein-RNA mixtures.^1–5^ Upon phase separation, these organelles emerge as droplets containing condensed component proteins, from the cytoplasm or nucleoplasm where the component proteins are dilute. Membraneless organelles have been associated with many cellular functions. In particular, the condensation of component proteins may facilitate the assembly of complexes for biochemical processes, as exemplified by the nucleolus, an organelle with a primary function in ribosome pre-assembly. Component proteins usually contain intrinsically disordered regions (IDRs) and RNA-binding domains. The effects of RNA on LLPS have been investigated in many experimental studies, but conflicting results have been reported.^2,6–15^ The aim of this study was to use computation to elucidate potential roles of RNA and other regulatory components in LLPS and the underlying physical mechanisms.

A hallmark of LLPS (in contrast to, e.g., aggregation) is thermodynamic reversibility. That is, droplets can easily dissolve upon raising temperature or salt concentration and can reform when conditions are reverted. The reversibility comes because the two phases, droplet and dispersed, are in thermodynamic equilibrium. The two phases separate when the molecules can achieve the same low free energy by adopting two distinct types of configurations, with disparate concentrations and extents of intermolecular bonding.^16^ The low free energy is achieved by distinct means in the two phases: e.g., high entropy (due to low concentration) in the dispersed phase but low enthalpy (from intermolecular bonding) in the droplet phase. Raising temperature or salt concentration leads to reduced stabilization of the droplet phase by intermolecular bonding. Beyond a critical point, the distinction between the two phases vanishes, and hence droplets dissolve. The thermodynamic equilibrium between the two phases is characterized by a phase diagram, or binodal, which is specified by the concentrations in the two coexisting phases for conditions below the critical point. In general, the stronger the intermolecular bonding, the higher the critical point.

Among the experimental studies on LLPS of protein-RNA mixtures, most reported that the addition of RNA promoted LLPS, indicated by either reduced threshold protein concentrations for droplet formation or increased critical salt concentrations.^2,7,8,10–13^ Reinforcing the promotional effects, both Saha et al.^12^ and Smith et al.^13^ found that molecules that competed for RNA binding with the droplet-forming proteins also suppressed phase separation. Intriguingly, Burke et al.,^6^ Zhang et al.,^9^ and Banerjee et al.^14^ observed that RNA promoted LLPS up to a point in the RNA to protein molar ratio; further increase in RNA led to LLPS suppression. The mechanism of such a dual effect remains unresolved; Zhang et al. suggested screening of interprotein electrostatic interactions by RNA (akin to salt) at higher concentrations, whereas Banerjee et al. speculated that increased RNA led to a charge inversion of the protein component. In yet another variation, Wei et al.^15^ recently found that RNA had little effects on the threshold protein concentration and critical salt concentration, but significantly reduced the protein concentration in the droplet phase. The binodals in the absence and presence of RNA were fitted to a polymer theory, with RNA treated only implicitly. Based on the fitting parameter values, the authors attributed the leftward shift of the high-concentration arm of the binodal in the presence of RNA to the latter’s weakening of interprotein attraction.

It should be recognized that the effects of RNA on protein LLPS are a reflection of intermolecular interactions rather than anything uniqueness about RNA. Indeed, similar promotional effects were observed in one case when RNA was replaced by heparin (which, like RNA, is an acidic polymer and thus presumably binds to the same sites on the droplet-forming protein)^10^ and in another case, instead of RNA, the regulatory component was arginine-rich peptides (expected to bind to acidic tracks, rather than RNA-binding sites, of the droplet-forming protein).^11^ LLPS also occurs for, and in fact, was first observed on, globular proteins.^17,18^ The phase separation of droplet-forming proteins was suppressed, with decreased critical temperatures, when another protein was added.^19,20^ For both disordered and globular proteins, LLPS was promoted by crowding agents such as polyethylene glycol (PEG) or Ficoll.^7,8,21,22^ Interestingly, in the latter study, bovine serum albumin (BSA) was found to suppress the phase separation of hnRNPA1 or its IDR fragment, but promoted, as if as a crowding agent, the phase separation of a protein-RNA mixture. Moreover, lysozyme exhibited a concentration-dependent dual effect on the LLPS of the IDR fragment of FUS (FUS_IDR_). In short, RNA and other regulatory components can exert similarly diverse effects on the LLPS of disordered and globular proteins. This is not surprising, since the LLPS of disordered and globular proteins have a common physical basis, though their phase behaviors show characteristic differences.^23^

Experimental studies have presented many observations on LLPS that beg for mechanistic answers. In principle, theoretical calculations and molecular simulations can provide these answers, but still face significant technical challenges. Polymer theories,^24,25^ coarse-grained simulations,^26^ and hybrids thereof^27^ have been used to determine phase diagrams of chain molecules, in particular as models of intrinsically disordered proteins (IDPs), and to address several important questions. These include how phase boundaries are affected by charge patterns along the sequence, chain length, charge mutations, and salt. The study that relates most closely to the present one is by Lin et al.,^25^ who treated the phase separation of a mixture of two IDP sequences within a polymer theory. All these approaches have limitations and hence their predictions are necessarily qualitative. In particular, polymer theories often provide only a very crude account of excluded-volume effects.^23^

Here we used Gibbs-ensemble simulations to investigate how regulatory components affect protein droplet formation, a pressing question that has received only scant attention in previous theoretical and simulation studies. While the promotion of LLPS by crowding agents can be understood from the perspective of excluded-volume effects,^28,29^ when RNA or another regulatory component is attractive toward the droplet-forming component and hence is recruited into the droplet phase, the physical rules governing the foregoing disparate effects on LLPS are yet to be established. Toward that aim, we chose a relatively simple model for protein-regulator mixtures, to calculate phase diagrams over a wide range of parameter values. To our knowledge, this is the first computational study devoted to uncovering the physical determinants underlying the roles of RNA and other regulatory components in LLPS. Our results qualitatively recapitulate the behaviors observed in the experimental studies, and provide a physical basis, in terms of the relative strengths of protein-protein and protein-regulator interactions, to reconcile the apparently conflicting effects of RNA on droplet formation.

## Results

### A simple model for protein-regulator mixtures

The droplet phase of protein-RNA mixtures, as true for LLPS in general, is stabilized by bonding networks of the molecules.^23^ As alluded to above, we reason that the disparate effects on LLPS are generic instead of unique to RNA, and hence can be captured by models that possess common features of mixtures of proteins with RNA or other regulatory components, in particular, polyvalent interactions. Here we used patchy particles to model protein and regulatory molecules (Figure 1a). These particles, with attractive patches decorated on a hard core, which in our case is spherical, have been used to model proteins in LLPS.^30–33^

**Figure 1.**
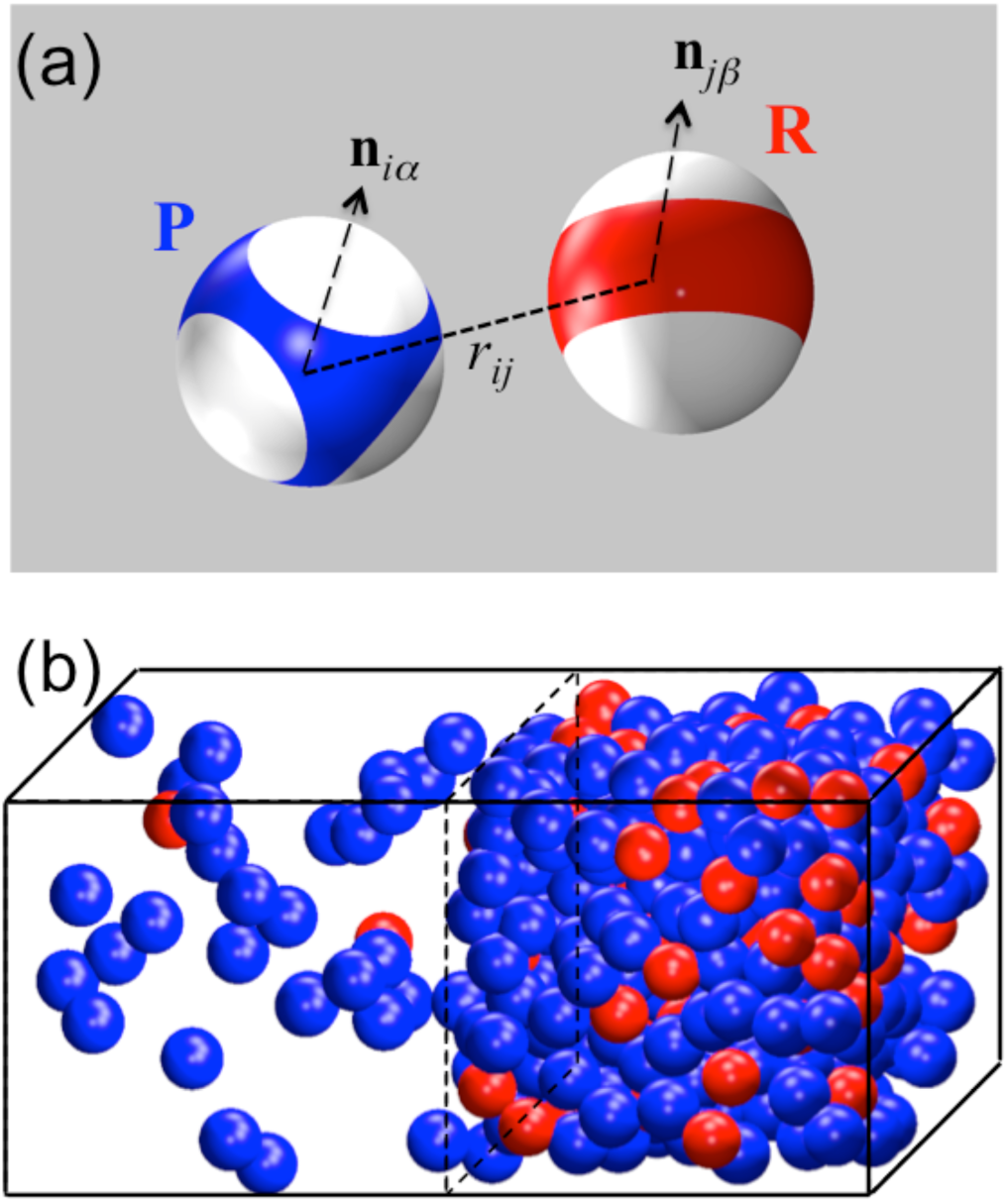
Illustration of the patchy particle model for protein-regulator mixtures and its two phases. **(a)** P and R particles, with cores in blue and red, respectively, and patches in white. **(b)** The low- and high-concentrations phases, after reaching equilibrium in a Gibbs ensemble Monte Carlo simulation. Patches are not displayed.

The details of the protein and regulator models were chosen based on a number of observations regarding LLPS of protein-regulator mixtures. First of all, in many experimental studies where LLPS has been demonstrated, there exists a single “driver” protein that can form droplets on its own, whereas RNA or other regulatory components can modulate the phase boundaries. Correspondingly, interprotein interactions have to be strongly attractive (for droplet formation), whereas inter-regulator interactions are weakly attractive or even purely repulsive. The latter scenario is certainly true for RNA, due to the uniform anionic charges of the phosphates. Meanwhile protein-regulator interactions are expected to be highly dependent on the particular molecules involved. Lastly, all particles occupy volume so they experience steric repulsion, which, much like the attractive interactions between patches, should play a role in determining phase diagrams, as demonstrated by effects of crowding agents.

In our models, protein-like (“P”) and regulatory (“R”) particles have 4 and 2 patches, respectively, to be denoted as *m*_S_, where S = P or R. The patches together cover the same fraction, *χ*, of the surface areas of the hard cores, which have the same diameter *σ* for both types of particles. Each patch is a spherical disk, with the spanning polar angle *θ*_S_ given by

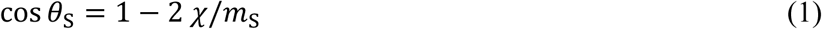

Each patch is identified by the unit vector from the particle center to the patch center, denoted by **n** with appropriate indices. The centers of the patches are positioned as a tetrahedron on a P particle but at the poles on an R particle. The greater spreading of the patches on a P particle gives it an advantage in forming bonds with multiple partner particles. A bond is formed when a patch on one particle is in contact with a patch on another particle.

The interaction energy between any two particles, *i* and *j*, has a directional square-well form,^34^

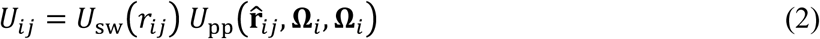

where *r*_*ij*_ is the distance between the particle centers, 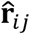 is the unit vector between them, **Ω**_*i*_ and **Ω**_*j*_ denote the orientations of the particles. The square-well part has the usual form,

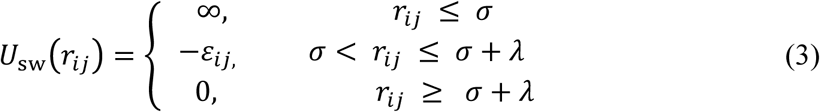

where *ε*_*ij*_ and *λ* are the strength and width, respectively, of the interparticle attraction. The interparticle attraction operates only when the patches of the two particles form a bond. The latter condition is specified by 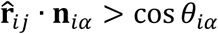 and 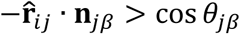, where *α* or *β* refers to any of the patches on particle *i* or *j*, and *θ*_*iα*_ or *θ*_*jβ*_ is the corresponding spanning polar angle of the patch. Hence

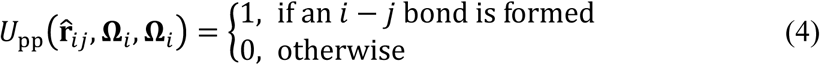

Note that each patch can simultaneously form bonds with multiple partner particles. The total energy of the two-component patchy particle system is the sum of all pairwise interactions between particles.

In the present work, *χ* was fixed at 0.7 and *λ* was fixed at 0.5*σ*. We set *ε*_RR_ = 0 (so the R particles experienced only steric repulsion between each other), and varied the strength of attraction between P and R particles, *ε*_PR_ = 0, over a wide range, from 0 to 1.5*ε*_PP_.

### Phase diagrams

For each *ε*_PR_, we used Gibbs-ensemble Monte Carlo simulations^35^ to determine phase diagrams for the protein-regulator model system over a wide range of molar ratios between P and R particles. In these simulations, fixed numbers of P and R particles were placed in a box with a fixed volume and maintained at a constant temperature. The box was divided into two regions, and the volume and the particle numbers in each region were changed to achieve equality in pressure and chemical potentials. After reaching equilibrium, the two regions represent different phases, with distinct concentrations (denoted by *ρ* with appropriate subscript and superscript) for the two types of particles (Figure 1b). We refer to the low- and high-concentration phases by I and II. A binodal consists of coexistence concentrations, 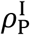 and 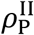, at a series of temperatures. For notational simplicity, hereafter all lengths and energies are, respectively, in units of the particle diameter *σ* and the strength of P-P attraction, *ε*_PP_ (equivalent to defining *σ* = 1 and *ε*_PP_ = 1). Likewise the temperature, *T*, is in units of *ε*_PP_/*k*_B_, where *k*_B_ is the Boltzmann constant. We denote the ratio of total R and P particle numbers as *X*.

The effects of the R particles on the phase diagrams differ qualitatively depending on *ε*_PR_ (Figure 2). They fall into three regimes, at low, moderate, and high *ε*_PR_ values. The low-*ε*_PR_ regime is represented by the results at *ε*_PR_ = 0 (Figure 2a). In this case, R particles promote LLPS. The critical temperature (*T*_c_) for the pure P system (i.e., *X* = 0) is 0.75 (in agreement with a previous study^34^). With increasing *X*, *T*_c_ increases (indicated by an upward gray arrow in Figure 2a) and the binodal broadens in both the low- and high-concentration arms, in a near symmetric fashion. Hence the threshold P concentration 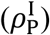 for phase separation, at a fixed temperature below *T*_c_ (e.g., *T* = 0.7, indicated by a horizontal line in Figure 2a), is a decreasing function of *X* (Figure 2a inset).

**Figure 2.**
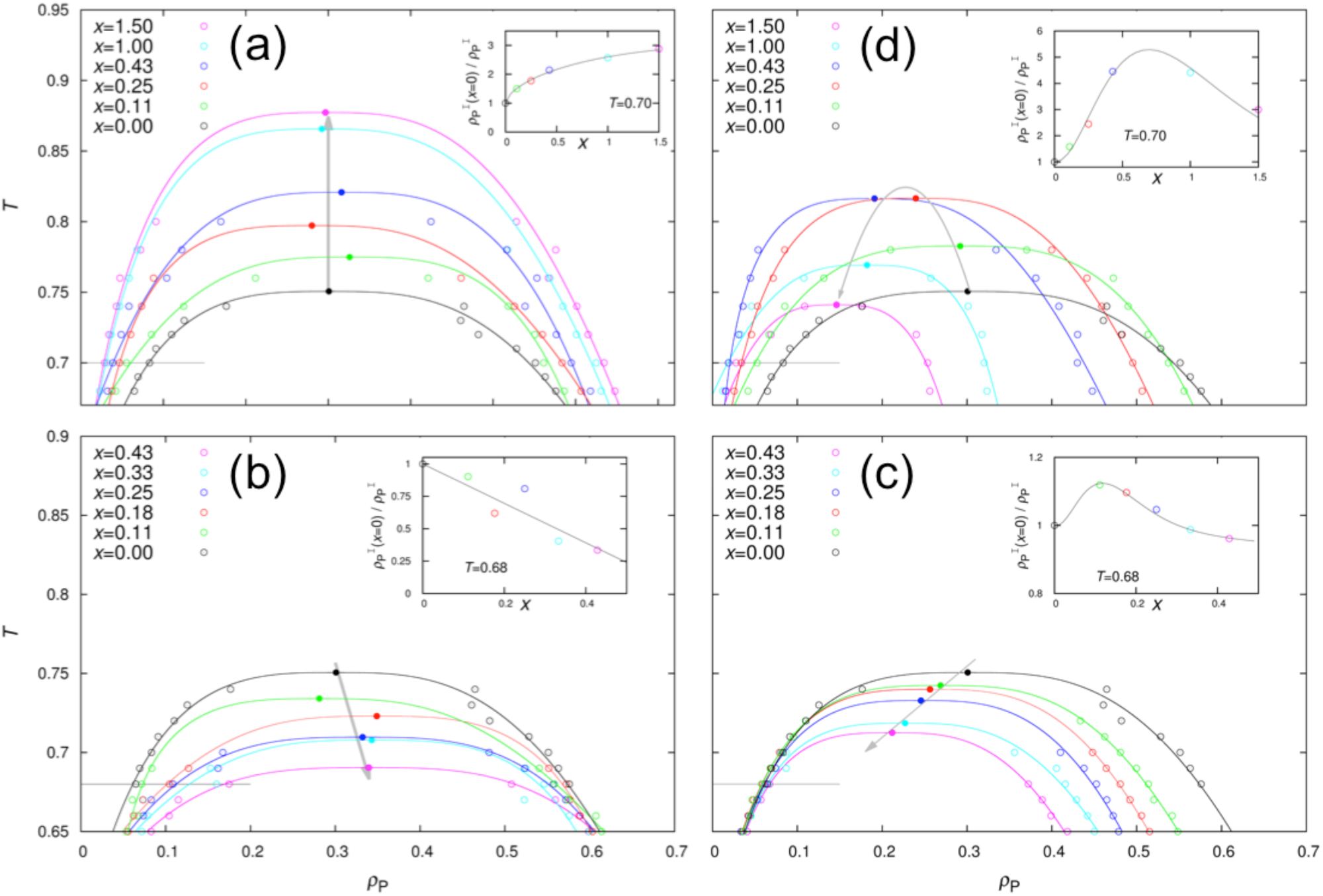
Phase diagrams. Results at *ε*_PR_ = 0.0, 0.5, 1.0, and 1.35 are shown in **(a)** – **(d)**, respectively. Open symbols are calculation results, and solid curves are fits to equation (7). The critical points from the fits are shown as filled symbols, with an arrowed gray line or parabola indicating the change with increasing *X*. A horizontal line through the left arm of the binodal indicates the temperature chosen for displaying results in the inset here and also in subsequent figures. Inset: ratio of threshold P concentrations between the pure P system and P-R mixture; a smooth curve is drawn to guide the eye.

As *ε*_PR_ increases from 0, the ability of the R particles deteriorates in promoting phase separation. At around *ε*_PR_ = 0.25, they begin to suppress phase separation. The suppressing effects are illustrated by the results at *ε*_PR_ = 0.5 (Figure 2b), which are essentially opposite to those at *ε*_PR_ = 0. Specifically, now *T*_c_ decreases monotonically with *X*, and the binodal narrows in both the low- and high-concentration arms, though more pronounced in the former than in the latter. With further increase in *ε*_PR_, the *T*_c_ decrease by the R particles is even somewhat greater at *ε*_PR_ = 0.75 (Supplementary Fig. S1a) but then becomes tempered at *ε*_PR_ = 1.0 (Figure 2c). Meanwhile, the effect on 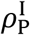 nearly vanishes, with a hint of a small decrease around *X* = 0.1 for *T* = 0.68 (Figure 2c inset). Instead the most prominent effect is on 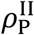, which decreases by 35% at *X* = 0.43 relative to the counterpart in the pure P system.

At even higher *ε*_PR_, a third regime emerges, as illustrated by the results at *ε*_PR_ = 1.35 (Figure 2d; see also Supplementary Fig. S1b for results at *ε*_PR_ = 1.25). Here the effects of the R particles depend on the R to P molar ratio. With increasing *X*, *T*_c_ initially increases, reaching a maximum at *X* = 0.35 (indicated by a parabolic gray arrow in Figure 2d). Then *T*_c_ gradually decreases, going below the counterpart for the pure P system at approximately *X* = 1.3. The threshold P concentration for phase separation has a similar turnover behavior, reaching a minimum at *X* = 0.7 for *T* = 0.7 (Figure 2d inset). In contrast, 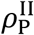 is a monotonically decreasing function of *X*. When *ε*_PR_ is further increased to 1.5, the maximal *T*_c_ shifts to a higher *X* value of approximately 0.65 and promotion of LLPS (i.e., *T*_c_ > the counterpart for the pure P system) persists over a wider range of *X* (Supplementary Fig. S1c).

### Partitioning of R particles in two phases

While Figure 2 displays the concentrations of the P particles in the two coexisting phases, the phase equilibrium is fully characterized by the concentrations of both types of particles. In Figure 3 we present the compositions of the two phases for the system with the four representative *ε*_PR_ values at a given temperature. At each *X*, the compositions are shown as a pair of points, 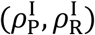 and 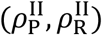, connected by a black line (known as a tie line).

**Figure 3.**
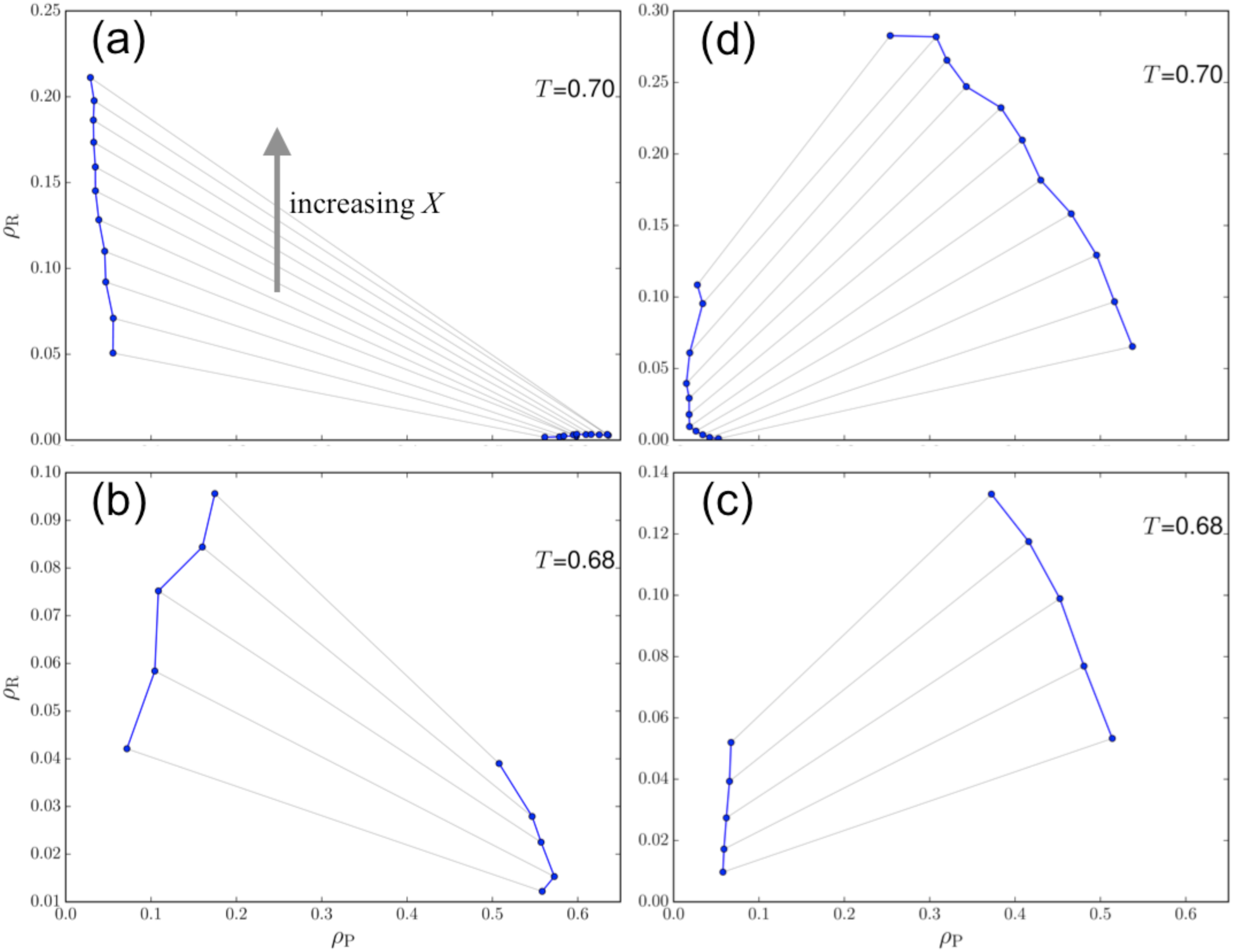
Partitioning of P and R particles in the two phases. Results at *ε*_PR_ = 0.0, 0.5, 1.0, and 1.35 are shown in **(a)** – **(d)**, respectively, at the temperatures indicated. A pair of points connected by a black line represent 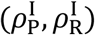 and 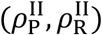 for the system at a given *X*.

At *ε*_PR_ = 0, the R concentrations in the droplet phase are miniscule for all *X* values, while those in the dispersed phase increase with increasing *X* (Figure 3a). These R particles preferentially partition in the dispersed phase, as expected for volume-exclusion crowders. In doing so, they displace the P particles into the droplet phase, as indicated by higher and higher 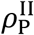 and lower and lower 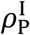 at increasing *X*. The latter is already seen in Figure 2a as near symmetric broadening of the binodal. Preferential partitioning in the dispersed phase occurs at *ε*_PR_ up to 0.2, where steric repulsion of R particles still dominates.

With increasing *ε*_PR_, more and more R particles are recruited into the droplet phase. Whereas the R particles still prefer the dispersed phase at *ε*_PR_ = 0.5, as indicated by tie lines with a negative slope (i.e., 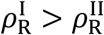; Figure 3b), the preference switches to the droplet phase at *ε*_PR_ = 1.0, where the tie lines now have a positive slope (Figure 3c). The preference of the R particles for the droplet phase further strengthens at *ε*_PR_ = 1.35 (Figure 3d). As more and more R particles are recruited into the droplet phase, more and more P particles are displaced from it. The latter trend is already seen in Figure 2c,d as significant leftward shifts by the high-concentration arm of the binodals. The simple reason for the displacement is that both P and R particles occupy volume so that the total concentration of particles that can be accommodated in the droplet phase is limited.

### Correlation between *T*_c_ and average number of bonds

The critical temperature for LLPS is determined by the strength of the bonding networks in the droplet phase. We measured the latter by the average number of bonds, *n*_b_, formed per particle at a given temperature. In Figure 4 we present *T*_c_ and *n*_b_ side by side, as functions of *X*. For all the four representative *ε*_PR_ values, the dependences of *T*_c_ on *X* qualitatively follow those of *n*_b_. Specifically, at *ε*_PR_ = 0, both *T*_c_ and *n*_b_ increase with increasing *X* (though a turnover may be expected at very high *X*; Figure 4a), whereas at *ε*_PR_ = 0.5 and 1.0 both *T*_c_ and *n*_b_ decrease monotonically with *X* (Figure 4b,c). At *ε*_PR_ = 1.35, both *T*_c_ and *n*_b_ exhibit a turnover, with the maxima occurring at nearly the same X value (around 0.3).

**Figure 4.**
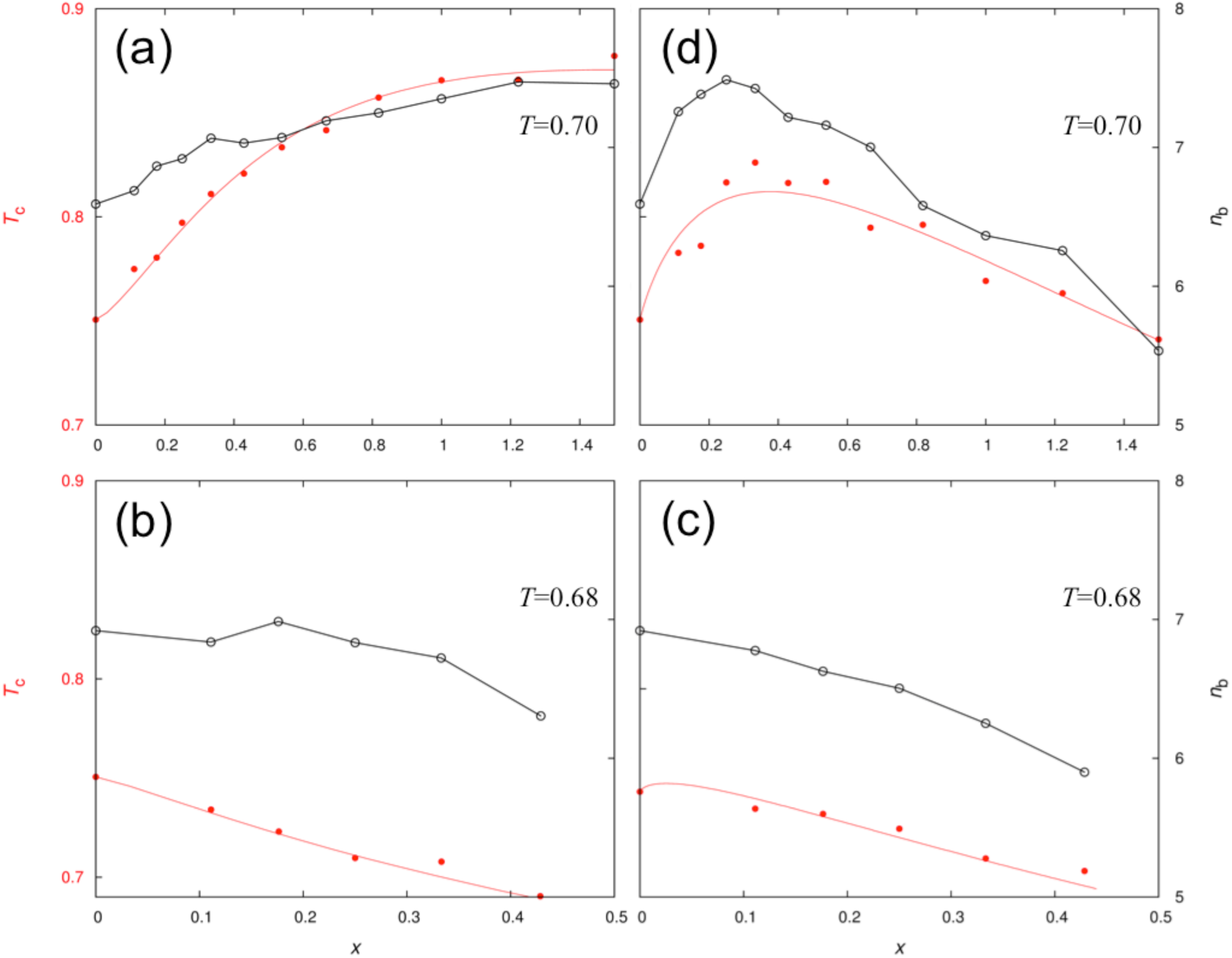
Correlation between critical temperature and average number of bonds per particle in the droplet phase. Results at *ε*_PR_ = 0.0, 0.5, 1.0, and 1.35 are shown in **(a)** – **(d)**, respectively. A smooth curve through the *T*_c_ data points is drawn to guide the eye; neighboring data points for *n*_b_ are connected by line segments. The latter are for the temperatures indicated.

### Physical mechanisms for diverse effects of R particles

We are now in a position to elucidate the physical mechanisms behind the various effects of the R particles on phase separation, by examining how they affect *n*_b_. At low *ε*_PR_, as typified by the situation at *ε*_PR_ = 0, the R particles preferentially partition into the dispersed phase, thereby displacing more and more of the P particles into the droplet phase. As 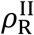 increases, each P particle is surrounded by more and more other P particles, and hence can form more bonds. This explains why *n*_b_ at *T* = 0.7 increases from 6.6 at *X* = 0 to 7.5 at *X* = 1.5 (Figure 4a). Therefore volume-exclusion crowders promote LLPS, as observed for both protein-RNA mixtures and globular proteins,^7,8,21,22^ by preferentially partitioning into the dispersed phase and thereby passively displacing droplet-forming proteins into the droplet phase to form stronger bonding networks.

To better explain how the R particles affect *n*_b_ at moderate and high *ε*_PR_, we further decomposed it. *n*_b_ is the average of *n*_b;P_, the number of bonds formed per P particle, and *n*^b;R^, the counterpart per R particle, weighted by their populations (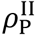 and 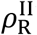):

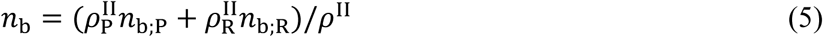

where 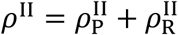 is the total concentration of particles in the droplet phase. Since a P particle can form bonds with both P and R partners (whereas an R particle can only have P partners), we further separated *n*_b;P_ by the types of partner particles,

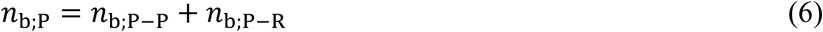

In Figure 5 we present the decomposition of *n*_b_ as well as 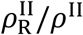 at *ε*_PR_ = 0.5, 1.0, and 1.35 for a given temperature. At *ε*_PR_ = 0.5, the R particles have lower bonding ability than the P particles (recall that *ε*_PP_ = 1 by definition). Indeed, when surrounded by pure P particles, a P on average forms 7 bonds [*n*_b;P_ at *X* = 0 in Figure 5a, at *T* = 0.68), while an R forms only 4.5 bonds (extrapolation of *n*_b;R_ to *X* = 0). As *X* increases, more and more R particles are recruited into the droplet phase, and hence they weight down the overall *n*_b_. In addition, at the highest *X* value (i.e., 0.43) studied, *n*_b;P_ is significantly lower (to 6.5 bonds) than the counterpart in the pure P system. The latter comes about because, due to the displacement of P particles by R particles, the loss in *n*_b;P-P_, the number of bonds with P partners, is not fully compensated by the gain in *n*_b;P-R_, the number of bonds with R partners.

**Figure 5.**
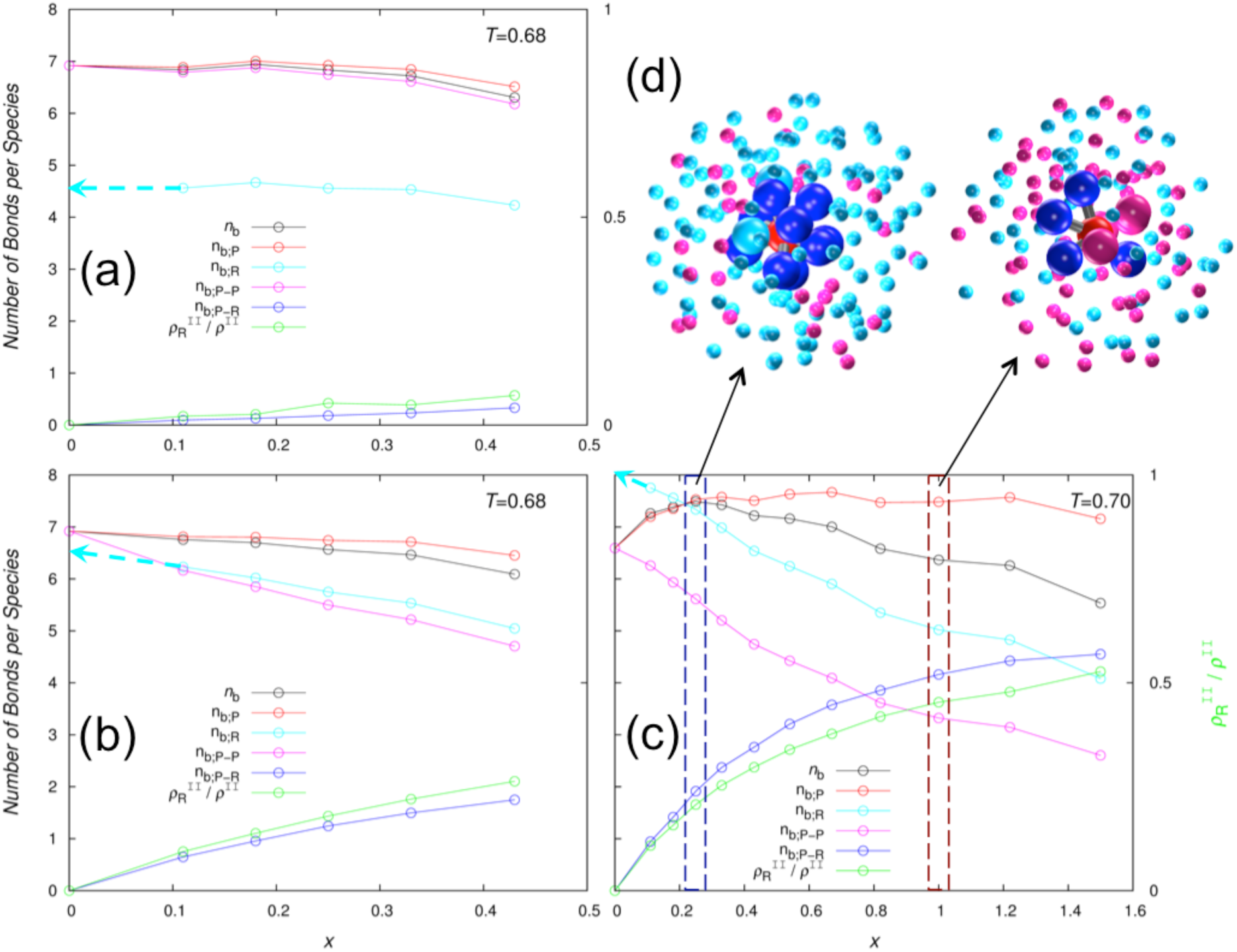
The average number of bonds per particle in the droplet phase and its decomposition by species. **(a)** – **(c)** displays bond information (left ordinate axis) and molar fraction of R particles in the droplet phase, 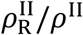, (right ordinate axis) at *ε*_PR_ = 0.5, 1.0, and 1.35, respectively, for the temperatures indicated. **(d)** Illustration of bonding networks in the droplet phase at *ε*_PR_ = 1.35, where R particles strengthen bonding at small *X* (left, *X* = 0.25) but disrupt bonding at large *X* (left, *X* = 1.0). In the central cluster, an R particle and its bonded P neighbors (9 in the left and 4 in the right) are shown as full-size spheres in red and blue, respectively, with bonds in gray; nonbonded P and R neighbors (2 Ps in the left and 4 Rs in the right) are shown as full-size spheres in cyan and magenta, respectively. The latter coloring scheme is also used for the more distant P and R particles, shown with radius reduced to 40%.

The behavior of *n*_b_ at *ε*_PR_ = 1.0 is similar to that at *ε*_PR_ = 0.5, but shows subtle differences (Figure 5b). The bonding ability of the R particles is now higher. Still, when surrounded by pure P particles, an R still lags behind a P in the number of bonds formed (6.5 for R versus 7 for P), due to less spreading of the patches on the R particles. Moreover, *n*_b;R_ now decreases significantly with increasing *X*, due to decreasing 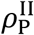 and hence a less number of bonding partners. The resulting widening gap between *n*_b;P_ and *n*_b;P_, coupled with the increasing R population, accounts for most of the decrease in *n*_b_ with increasing *X*. Lastly, *n*_b;R_ itself decreases with increasing *X*, again because the gain in *n*_b;P-P_ does not keep pace with the loss in *n*_b;P-P_. In short, at moderate *ε*_RP_, the R particles, by partitioning into the droplet phase and displacing P particles, actively weaken bonding networks there and thereby suppress LLPS.

At *ε*_PR_ = 1.35, the stronger R-P attraction (relative to the P-P attraction) enables an R to form more bonds than a P when each is surrounded by pure P particles (8 for R versus 6.5 for P at *T* = 0.7; Figure 5c). With the enhanced bonding ability for R, as *X* increases from 0 and more R particles displace P particles, the gain in bonds for a P formed with R particles is greater than the loss in bonds with other P particles (Figure 5d, left). Consequently, *n*_b;P_, the average number of bonds formed by a P, initially increases with increasing *X*. After *X* = 0.25, *n*_b;P_ levels off. Meanwhile *n*_b;R_ decreases monotonically with *X* due to the displacement of partners (i.e., P particles) by R particles (Figure 5d, right). At around *X* = 0.3, the weighted average of *n*_b;P_ and *n*_b;R_, i.e., *n*_b_, thus reaches maximum, and hence here the R particles have optimal ability in promoting LLPS. As *X* increases further and further and more and more P particles are displaced from the droplet phase, the R particles cannot form enough bonds so eventually they suppress phase separation.

## Discussion

We have calculated the phase diagrams of a model protein-regulator system over wide ranges of parameter values, in order to elucidate potential roles of RNA and other regulatory components in LLPS and the underlying physical mechanisms. The calculation results demonstrate that regulatory components can exhibit diverse effects on phase separation, depending on the relative strengths of protein-protein and protein-regulator interactions and the regulator to protein molar ratio (Figure 6). If the regulator is at most only weakly attractive toward the protein, it behaves like a volume-exclusion crowder in promoting LLPS, with increasing effects at higher regulator to protein molar ratios. (This trend may stop and even reverse eventually when the regulator is forced into the droplet phase and disrupts bonding networks there). In contrast, with moderate attraction toward the protein, the regulator suppresses LLPS (“active suppressor”), with effects monotonically increasing with the regulator to protein molar ratio. With strong protein-regulator interactions, the regulator initially promotes LLPS (“active promoter”), reaching a maximal effect at a certain regulator to protein molar ratio. But at higher and higher amounts of the regulator, the promotional effect subsides and eventually turns into suppression of LLPS.

**Figure 6.**
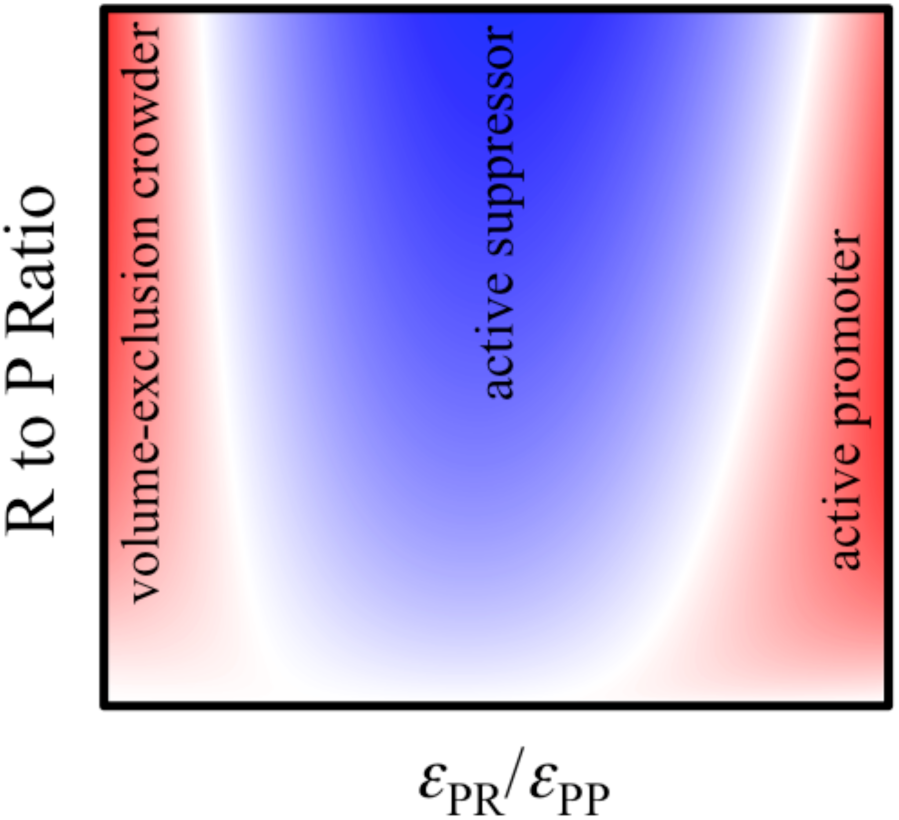
Expected effects of regulatory (“R”) particles on the droplet formation of protein (“P”) particles. Suppression of LLPS occurs at moderate P-R attraction (blue color), whereas promotion of LLPS occurs at either very weak or strong P-R attraction (red color). The promotion can turn into suppression of LLPS at high R to P ratios.

One obvious limitation of our model is that the protein and regulator are treated as rigid, whereas proteins forming membraneless organelles usually contain IDRs, and regulatory RNA can be flexible molecules as well. We contend, as further discussed below, our findings and the resulting physical insights transcend the divide between rigid and flexible molecules. Other recent theoretical and simulation studies have modeled flexible chain molecules.^24–27^ These studies have not addressed the central question posed here, i.e., the effects of regulatory components on protein LLPS. It will be interesting to extend these studies to test the applicability of our conclusions on flexible molecules.

To elucidate the underlying physical mechanisms of the effects of regulatory components, we have found it useful to characterize bonding networks in the droplet phase, specifically, by the average number of bonds per particle and its decomposition by species. In general, molecular interaction networks, or transient clusters, are crucial to the physical understanding of dense macromolecular systems, such as those in cellular environments.^16,36^

Our results qualitatively recapitulate the behaviors observed in experimental studies of protein-crowder, protein-protein, and protein-RNA mixtures, and provide a physical basis to reconcile conflicting reported results for the effects of RNA on protein droplet formation. Crowding agents such as PEG and Ficoll are found to promote LLPS for protein-RNA mixtures and globular proteins.^7,8,21,22^ Our calculation results with low *ε*_PR_ show that the promotional effect of such crowders comes because they preferentially partition in the dispersed phase, thereby passively displacing droplet-forming proteins into the droplet phase to strengthen the bonding networks there.

The critical temperature for LLPS of *γ*D crystallin decreased upon adding βB1 crystallin.^19^ Similarly, droplet formation of a monoclonal antibody and hnRNPA1, respectively, was suppressed by human serum albumin^20^ and BSA.^22^ The changes in *T*_c_ and the tie lines of the protein mixtures in the first two studies are similar to our results obtained at moderate *ε*_PR_. In this case, the regulatory protein has weaker attraction toward the droplet-forming protein than the latter toward itself. Still, the regulatory protein is recruited to the droplet phase, and thereby displaces the droplet-forming protein and weakens the bonding networks there. Both the recruitment and displacement features identified by our study were directly visualized by fluorescence microscopy for the hnRNPA1-BSA systems, whereas PEG and Ficoll, also as expected, were excluded from the droplet phase.^22^ Displacement occurs simple because there is a limit to the total concentration of macromolecules in the droplet phase.

Promotional effects of RNA on LLPS have been reported in a number of studies.^2,7,8,10–13^ We explain these results by suggesting that the RNA molecules in these studies have stronger attraction toward the droplet-forming proteins than the latter toward themselves, and therefore actively strengthen bonding networks in the droplet phase. This suggestion is quite reasonable, since the droplet-forming proteins all contain specific RNA-binding domains to achieve strong bonding with RNA. We further hypothesize that the promotional effects in these studies were obtained when the amounts of RNA added were sufficiently low, and predict that adding higher amounts of RNA would eventually lead to LLPS suppression. Indeed, such dual effects have been observed in three studies.^6,9,14^ The present study reveals that the suppression at high RNA-to-protein molar ratios arises from the displacement of proteins by RNA from the droplet phase, leading to an insufficient level of proteins to maintain the strength of the bonding networks of the protein and RNA molecules. This mechanism is distinct from those by Zhang et al.^9^ and Banerjee et al.^14^, who speculated that RNA would either screen interprotein electrostatic interactions like diffuse salt ions or invert the protein net charge by strong binding to the protein surface. While these specific effects of RNA may exist, our mechanism is generally applicable to RNA and any other macromolecular regulators.

As support for our contention that the findings in the present study are applicable to both rigid and flexible molecules, a concentration-dependent dual effect was also found for lysozyme, a globular protein, on the LLPS of FUS_IDR_.^22^ In comparison, BSA did not show any sign of promoting the FUS_IDR_ LLPS. We can now explain the different effects of lysozyme and BSA in terms of their relative strengths of attraction toward FUS_IDR_. That is, we predict that FUS_IDR_ is more strongly attracted to lysozyme than to BSA. Interestingly, a recent small-angle neutron scattering study indicated exactly such differential strengths of attraction of the latter two proteins toward FlgM, an intrinsically disordered protein.^37^ While two regulatory proteins (lysozyme and BSA) can have divergent effects (active promoter vs. active suppressor) on the phase separation of a given protein (FUS_IDR_), a given regulatory protein can also have distinct effects on the phase separation of two different proteins. The latter was observed with BSA acting as an active suppressor for the LLPS of hnRNPA1 but as a volume-exclusion crowder for the LLPS of a protein-RNA mixture.^22^ Again, the distinct effects can be attributed to the differential strengths of attraction of the regulatory protein toward the droplet-forming proteins. We note that the strengths of intermolecular interactions can be directly measured by the second virial coefficient or the binding affinity, determined at concentrations below the threshold for LLPS.^13,15^

In another intriguing report,^15^ polyadenylate RNA had little effects on the threshold LAF-1 concentration for LLPS and critical salt concentration, but significantly reduced the protein concentration in the droplet phase. These effects are similar to our calculation results at *ε*_PR_ = 1.0, where the dominant effect of the R particles is to reduce 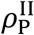 (Figure 2c). This comes about because most of these R particles partition into the droplet phase and displace P particles, thereby reducing 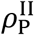 while largely preserving the strength of the bonding networks (Figure 5b). In contrast, Wei et al.^15^ related the reduction of 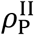 to weakening of interprotein attraction by RNA. That RNA would displace proteins in the droplet phase was not considered in the model used for their data fitting.

Our study has identified the relative strengths of protein-protein and protein-regulator interactions as an important physical determinant for the effects of regulatory components on protein droplet formation. In particular, active promotion of LLPS occurs only when RNA or other regulators have stronger attraction toward proteins than the latter toward themselves. Conversely, RNA is expected to behave like volume-exclusion crowders when droplet-forming proteins do not have much affinity for RNA. This can happen when droplet-forming proteins are deleted of their RNA-binding regions. Burke et al.^6^ actually tested such a construct, FUS_IDR_, for RNA effects and the results were null. It is possible that the FUS_IDR_ concentration in that test was too low, and that a higher FUS_IDR_ concentration may exist where LLPS can be induced by adding RNA. We hope that these and other predicted trends in Figure 6 will motivate experimental tests.

Different membraneless organelles may use RNA for different roles, some as clients whereas others for active promotion of LLPS. Our study demonstrates that these different roles can be fulfilled by modulating the relative strengths of protein-protein and protein-RNA interactions. Furthermore, if RNA does promote LLPS, it can do so only within a certain concentration range. This turnover behavior affords cells the ability to use RNA expression level for regulating the formation of membraneless organelles.

## Methods

The phase diagrams of the protein-regulator model system for *ε*_PR_ (strength of P-R attraction) between 0 and 1.5 and *X* (R to P molar ratio) between 0 and 1.5 were determined from Gibbs-ensemble Monte Carlo simulations.^35^ In most simulations, the total number of particles was 512 and the total concentration, *ρ*_0_, was 0.3, a setup where the binodal for the pure P system has been obtained previously;^34^ our simulations at *X* = 0 reproduced this result. Some simulations were also carried out with a total of 1000 particles (while preserving *ρ*_0_); the resulting coexistence concentrations were unchanged. In other simulations *ρ*_0_ was changed to 0.1 and the results are qualitatively consistent with those at *ρ*_0_ = 0.3. Each binodal was obtained from simulations at a series of temperature ranging from 0.65 to 0.80. At each temperature, the first 1 to 1.5 million cycles were discarded to ensure equilibration, and the subsequent 2 to 3 million cycles were used for data collection. Each cycle consisted of 500 attempts of particle displacement, 500 attempts of particle exchange, and 5 attempts of volume exchange.

The critical point for each binodal was determined by fitting the temperature dependence of the coexisting P concentrations, 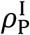 and 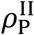, to the law of rectilinear diameter^38^

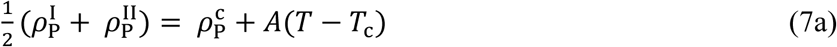

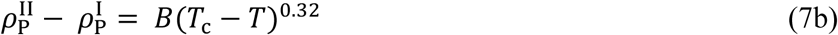

where 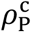 is the critical protein concentration, and *A* and *B* are other fitting parameters. Each fit was performed first by including data at all available temperatures and then gradually removing data that were closer to *T*_c_, until the fitted values of 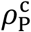 and *T*_c_ were stable.

## Data availability

The datasets generated and analyzed during the current study are available from the corresponding author on reasonable request.

## Acknowledgements

This work was supported by National Institutes of Health Grant GM118091 and US-Israel Binational Science Foundation Grant 2015376.

## Author Contributions

V.N. and H.-X.Z. designed the research. V.N. performed the research. V.N. and H.-X.Z. wrote the manuscript.

## Additional Information

### Supplementary information

accompanies this paper

### Competing Interests

The authors declare that they have no competing interests.

